# Regional patterns of neurodegeneration in a mouse model of proteinopathy

**DOI:** 10.1101/2025.08.17.670564

**Authors:** Dmytro Shepilov, Ibrahim Ahmed Khan, Hsiao-Jou Cortina Chen, Christine Rowley, Edward C. Harding, Florian T. Merkle

**Affiliations:** Institute of Metabolic Science, University of Cambridge, Cambridge, CB2 0QQ, UK; Department of Pharmacology, University of Cambridge, Cambridge, CB2 1QR, UK; Wellcome-MRC Cambridge Stem Cell Institute, University of Cambridge, Cambridge, CB2 0AW, UK

**Keywords:** scrapie, prion, neurodegeneration, formic acid, glia, neurons, ER stress, spongiosis

## Abstract

The aggregation of misfolded proteins is a hallmark of many neurodegenerative diseases, suggesting shared pathological mechanisms. However, the pathways by which protein misfolding in these proteinopathies lead to neuronal death remain unclear. Proteinopathies can be modelled in transgenic animals by expressing disease-causing mutations that promote protein aggregation, or in wild-type animals by injecting misfolded proteins (e.g. RML scrapie) that spread in a prion-like manner and recapitulate key neurodegenerative features, including gliosis, ER stress, and neuronal loss. Here, we map region-specific histopathological features of scrapie-induced neurodegeneration in the hippocampus, thalamus, cortex, and cerebellum during early (12 weeks post-inoculation) and late (20 weeks) stages of disease. Using a streamlined time-efficient protocol, we achieve reproducible paired sample collection and high-quality immunohistochemistry that is compatible with best practice in decontamination and containment. We found that among the tested markers of early pathology, thalamic astrocytic activation and spongiform degeneration were the most sensitive. By the late stage, there was widespread upregulation of IBA1^+^ microglia and GFAP^+^ astrocytes, accompanied by strong immunoreactivity of lysosomal marker LAMP1. LAMP1 expression in healthy brains was largely neuronal, but by 20 weeks it was significantly upregulated in astrocytes, suggesting their involvement in lysosomal pathology. The ER stress marker p-PERK was elevated in CA1/CA3 pyramidal neurons but minimal in the thalamus and cerebellum, where neuronal loss was most pronounced, suggesting region-specific mechanisms of degeneration. Overall, the thalamus and hippocampal CA1/CA3 areas exhibited the greatest pathological burden. Our shorter time-course, new pathological insights and safe handling protocols, and improved welfare, supports broader adoption of the RML scrapie model for resource-efficient studies of neurodegeneration and its prevention.

## INTRODUCTION

Neurodegenerative conditions like Alzheimer’s Disease and Related Dementias (ADRD) are large, unmet clinical needs that are growing rapidly as populations age (Livingston et al. 2024). These conditions are characterised by the accumulation of misfolded proteins that appear to propagate across the brain, often forming insoluble fibres and plaques associated with inflammation and neuronal loss. Though protein aggregates are common to many neurodegenerative diseases, they arise from diverse genetic and environmental causes. These include β-amyloid (Aβ) plaques and Tau tangles in Alzheimer’s Disease (AD) (Busche and Hyman 2020), Tau and TDP-43 aggregation in Frontotemporal Dementia (FTD) (Goedert et al. 2021), α-synuclein aggregation in Parkinson’s Disease (PD) and Lewy Body Dementia (LBD) (Ayers et al. 2022), and the aggregation of misfolded PrP in prion diseases such as Creutzfeldt-Jakob Disease (CJD) (Jones and Mead 2020). These neurodegenerative diseases can be considered proteinopathies as they are characterised by protein misfolding, aggregation, apparent spreading, and associated neuroinflammation and neuronal loss (Bayer 2015; Jaunmuktane and Brandner 2020; Mallucci, Klenerman, and Rubinsztein 2020; D. M. Wilson 3rd et al. 2023).

There is a lack of consensus about how best to study human proteinopathies in animal models to understand disease mechanisms and to develop new therapies. One approach is to model rare, highly-penetrant, disease-associated mutations in transgenic mice, such as PSEN1/2 and APP in mouse models of ADRD (Zhong et al. 2024; Qian, Li, and Ye 2024). Each model recapitulates part of the clinical presentation of sporadic forms of disease in humans, such as the accumulation of Aβ plaques (Sasaguri et al. 2022), and these have aided in the development of treatments such as the anti-Aβ antibody donanemab (Sims et al. 2023). However, other clinical features may be lacking, such as tau tangles seen in the most common form of late-onset Alzheimer’s disease known as LOAD (Jankowsky and Zheng 2017). There are still limited therapeutic options that target proteinopathy-related neurodegenerative processes downstream of Aβ or tau (Cummings et al. 2024). An alternative approach is to study ‘pure’ proteinopathy models to reveal shared neurodegenerative mechanisms that could be targeted to develop cross-cutting neuroprotective therapies. Such models should ideally show rapid and stereotyped neurodegenerative disease progression that recapitulates key features of human neurodegenerative disease including bona fide neuronal loss in brain regions relevant to human disease, be inducible across different time points and genetic backgrounds, and be readily accessible to groups around the world. One mouse model that meets these criteria is the Rocky Mountain Laboratory (RML) scrapie model of prion disease, in which brain homogenate from infected mice is injected into the brains of healthy mice to induce proteinopathy and neurodegeneration (Prusiner 1982; Freeman and Mallucci 2016).

While sporadic proteinopathies of accumulating misfolded PrP are rare, the conversion of endogenous PrP^C^ to misfolded PrP^Sc^, and its propagation across the brain and resulting neurodegeneration potently induce an unfolded protein response leading to both translational arrest and disruption in lysosomal trafficking and vacuolisation (spongiosis) that gives the brain a sponge-like appearance at late stages of disease (Freeman and Mallucci 2016; Lakkaraju et al. 2021). As disease progresses, RML scrapie mice show synaptic and neuronal loss, clear motor signs, and broad microglial activation and astrocytic reactivity. These molecular and clinical outcomes occur on a practical timescale of months for wild-type animals, require neither complex breeding nor long-term ageing paradigms (Halliday, Radford, and Mallucci 2014), and are useful for pre-clinical drug testing. Thus, the inoculation of PrP^sc^ represents a simple and tractable model to recapitulate naturally occurring neurodegenerative processes resulting from accumulating misfolded protein.

Specifically, this model can efficiently test for neuroprotective compounds that mitigate neuronal loss, glial activation, and the unfolded protein response and result in prolonged survival and reduced clinical signs and spongiosis pathology (Sidhom et al. 2022). Drugs that are neuroprotective in scrapie include the PERK inhibitor GSK2660414 that prevents PERK pathway activation, the integrated stress response inhibitor (ISRIB) that reverses the effect of eIF2α phosphorylation (Sidrauski et al. 2015), and the eIF2α phosphatase salubrinal (Boyce et al. 2005; Sidhom et al. 2022). The effort to repurpose well characterised clinically safe drugs for endoplasmatic reticulum (ER) stress inhibition culminated in the identification of trazodone, a commonly used antidepressant, which is also able to suppress eIF2α phosphorylation and restore translation (Halliday et al. 2017).

Despite the many advantages of scrapie models of proteinopathy, transgenic models are used much more widely (Zhong et al. 2024; Watts and Prusiner 2014). This underutilisation is likely driven by concerns over potential infectivity and safe handling of scrapie-infected material, which persists despite the fact that there are no known cases of scrapie infecting humans (Peden, Suleiman, and Barria 2021) and no epidemiological evidence of risks associated with scrapie in the food chain, which was first identified more than 200 years ago (Cann 2016; Houston and Andréoletti 2019). To render scrapie non-infectious, chemical treatments such as concentrated formic acid have been used for decades to decontaminate prion samples of diverse origins, including human and mouse brain tissue (Brown, Wolff, and Gajdusek 1990; Tateishi, Tashima, and Kitamoto 1991). Concentrated formic acid disaggregates beta-sheet rich structures such as the PrP^sc^ by disrupting hydrogen bonding, and can induce internal covalent modifications and denaturation in otherwise resistant proteins (W. E. Klunk and Pettegrew 1990; William E. Klunk, Xu, and Pettegrew 1994). When combined with fixation by paraformaldehyde, concentrated formic acid reduces scrapie infectivity from brain tissue by 5 to 8 log units as assessed by RT-QuiC seeding or bioassay. Using the inoculation of treated samples directly into the mouse brain as the gold standard bioassay for infectivity, this treatment almost completely abolishes infectivity (Taylor et al. 1997; Dong et al. 2021). A decontamination pipeline should facilitate flexible tissue processing for immunohistochemistry to gain new cellular and subcellular mechanistic insights over the time course of RML scrapie disease progression.

Here we describe streamlined protocols for the reproducible and scalable dissection of paired samples from RML scrapie-infected mice for pre-clinical studies of neurodegenerative processes. We provide a spatial and temporal map of scrapie disease progression and reveal new information about region-specific vulnerability to misfolded protein aggregation, such as in the thalamus. Our findings suggest a unique role of astrocytes in spongiosis spatially restricted patterns of the ER stress marker p-PERK, and their respective relations to neurodegeneration. These approaches are suitable for large numbers of paired samples required for well-powered pre-clinical drug testing and multiomic analysis.

## RESULTS

Despite its many advantages, RML scrapie is not yet a widely-used model of neurodegeneration. We set out to integrate safe practice with cellular and molecular characterisation of region-specific disease progression in an easy-to-use protocol that would make adoption appealing to a wider range of research labs. Here we describe a pipeline for studying neurodegenerative processes in wild-type mice inoculated with the RML scrapie until 20-wks post-inoculation. The method allows us to preserve paired-samples, on an efficient time course, while using best-practice decontamination procedures to prepare brain hemispheres for immunohistochemistry (IHC). We demonstrate the utility of these methods by building a spatial and temporal map of RML scrapie progression to generate cellular and molecular insights into disease.

### Streamlined short time course model of prion disease to study neurodegeneration

To establish this pipeline, we first refined an efficient dissection and preservation routine for animal experiments using blood withdrawal under terminal anaesthesia. This approach both preserves terminal blood that is normally lost during perfusion for biochemical analysis, and removes red blood cells (RBCs) from the brain to reduce autofluorescence during subsequent brain IHC. We then systematically dissected tissue components for either IHC or paraffin embedding, molecular biology/biochemistry, or omics methods (**Figure 1A).** We collected tissue at fixed time points of 12- and 20-weeks post RML inoculation, these are significantly shorter than traditional survival experiments facilitating direct timepoint comparison of histology at two key points in disease trajectory: 12-wk precedes the appearance of behavioural changes but molecular changes may be apparent, and since by 20-wk strong molecular and behavioural changes are apparent but most animals have not entered terminal stages of disease (Harding et al. 2024). To facilitate paired comparisons, we split dissected brains into two hemispheres, retaining one for overnight fixation in 4% paraformaldehyde (PFA), and the second for microdissection into different brain regions that were snap frozen alongside organs, plasma, and RBCs (**Figure 1B**).

**Figure 1.**
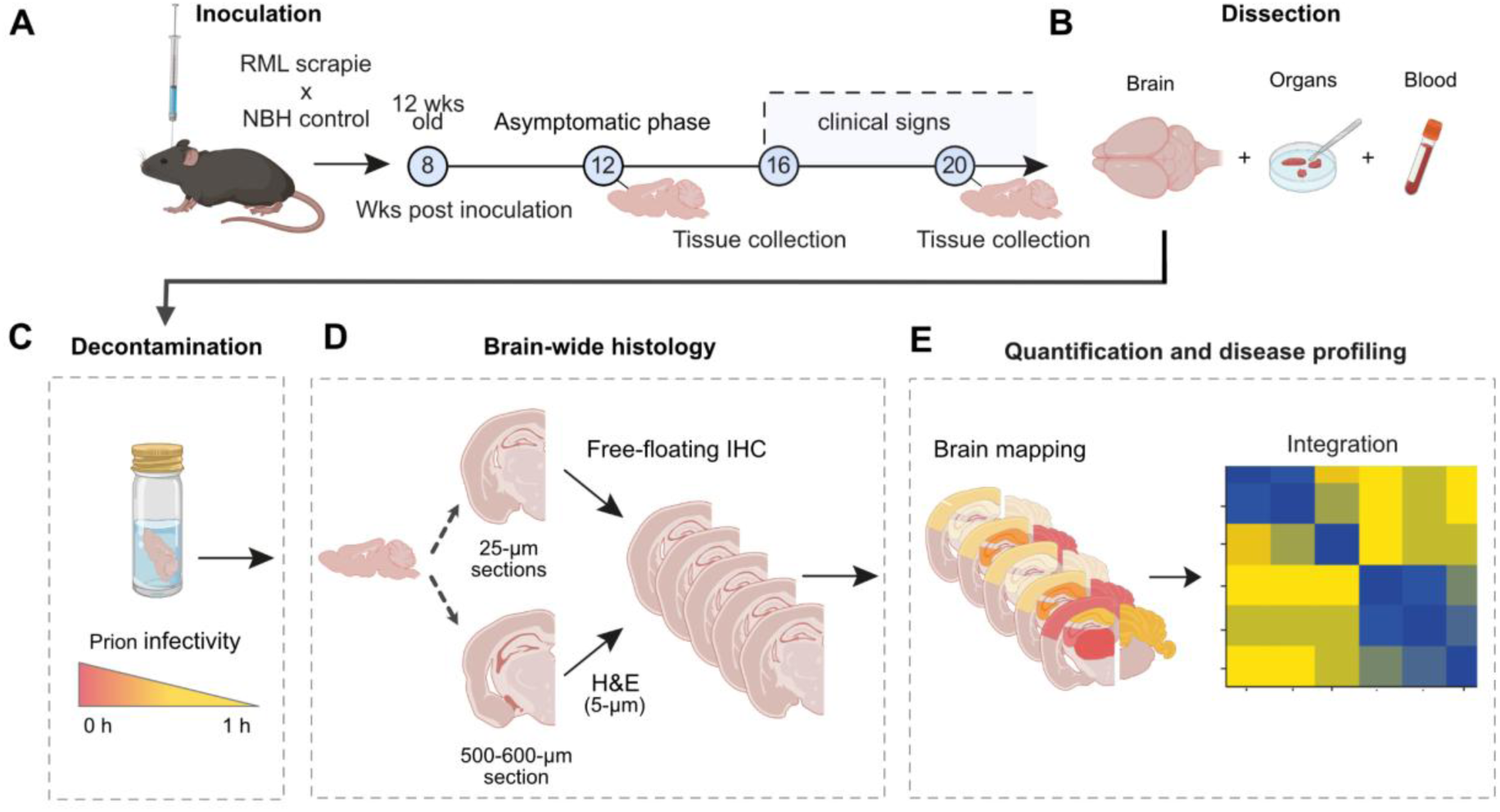
Schematic of histopathological mapping of RML scrapie-induced neurodegeneration in the mouse brain. **A)** Prion disease was induced in 4-6-week-old C57BL/6J mice by intracerebral inoculation with brain homogenate containing Rocky Mountain Laboratory (RML) scrapie prions, or normal brain homogenate (NBH) as a control. Tissue was collected at two time points during disease progression, corresponding either to the early asymptomatic phase (12 weeks post-inoculation; 12-wk) or after the appearance of clinical signs (20-wk) but well before terminal disease. **B)** Blood was collected by intracardial withdrawal, separated and stored as RBC and plasma prior to organs and brain collection to permit multiple types of analyses from paired samples. **C,D)** PFA-fixed hemispheres for histology were decontaminated in >95% formic acid for 1 h (C) and sectioned on a vibratome to obtain 25 -µm free-floating sections for immunohistochemistry while thick vibratome slices from the same brains were subsequently paraffin-embedded and sectioned on a microtome for H&E staining (D). **E)** Histopathological biomarkers were quantified from microscopic images and used to generate regional brain maps of prion disease and perform integrative analysis of morphological data.

Since PFA-fixed mouse brain hemispheres have a maximal thickness of 4-5 mm, we next tested whether immersing the hemisphere in >95% formic acid for 1 hour would facilitate downstream IHC (**Figure 1C**). Previous studies have shown that >95% formic acid treatment renders prion tissue up to 5 mm thick almost completely non-infectious (Brown, Wolff, and Gajdusek 1990; Tateishi, Tashima, and Kitamoto 1991; Taylor et al. 1997). We found that formic acid-treated hemispheres could be readily sectioned by vibratome and immunostained for most antigens, with simple but important adaptations as described below. Thick brain sections could also be embedded in paraffin for thin microtome sectioning and hematoxylin and eosin (H&E) staining (**Figure 1D**). Upon optimising IHC methods, we immunostained inoculated mouse brain sections and quantified immunoreactivity across key brain regions. This allowed us to create comprehensive brain maps of histopathological markers relevant to prion disease pathogenesis in the RML-inoculated mouse brain and integrate this data to model disease progression (**Figure 1E**).

### Improved immunofluorescence in formic acid-treated scrapie-inoculated brains

Formic acid nearly completely denatures proteins at room temperature when given at concentrations of ∼6M (Christensen et al. 2020), so we wondered how treatment of brains with >95% formic acid (>24M) would affect both tissue autofluorescence artefacts and immunoreactivity. To address this question, we first tested how treating control-injected brain tissue with >95% formic acid for 1 hour followed by washing with phosphate buffered saline (PBS) for 10 minutes or 24 hours would affect tissue integrity. We found that formic acid treatment increased brain hemisphere size, likely driven by osmosis, but hemispheres returned to normal size after 24 hours of washing in PBS (**Figure S1A-C**). We found that decontaminated brains retained enough structural integrity to enable coronal vibratome sectioning at 25 µm. When we imaged sections of formic acid-treated and untreated brains, we observed statistically significant increase in autofluorescence (**Figure S1D**) that we quantified across 405, 488, and 561 nm wavelengths of illumination (**Figure S1E-G**).

To mitigate the effects of decontamination-induced autofluorescence and to improve staining quality more generally, we considered several potential contributors to autofluorescence in brain tissue at each of these wavelengths including tryptophan (Em: 320), extracellular matrix elastin (Em: 420-520 nm), intracellular flavin adenine dinucleotide (FAD, Em: 520-560 nm), and reduced nicotinamide adenine dinucleotide (NADPH, Em: 520-560 nm). We next considered methods that could be applied reproducibly and in parallel to quench this autofluorescence, including incubation of free-floating sections with copper (II) sulfate (CuSO_4_) ^(^_Erben et al. 2016_^)^, the lipophilic diazo dye Sudan Black B (SBB) (Erben et al. 2016; Oliveira et al. 2010), or pre-bleaching by overnight LED-based illumination (Duong and Han 2013) (**Figure S1H-J)**. We found that both SBB and LED methods significantly (p<0.05) reduced autofluorescence intensity across 405, 488 and 561 nm, but CuSO_4_ treatment did not (**Figure S1K-N**).

We then immunostained mouse coronal sections from RML and NBH-inoculated mice (**Figure 2A, S2A**) with antibodies against anti-ionised calcium-binding adaptor molecule 1 (IBA1) and glial fibrillar acidic protein (GFAP) that are commonly used to identify microglia and astrocytes, respectively and focused on regions of interest such as the hippocampus (**Figure 2B**). We first confirmed these antigens were retained after formic acid treatment, and that the reduced background in tissues treated with SBB or by LED (**Figure S2A**) led to improved clarity upon imaging when imaged at 488 nm. This was evidenced by a greater signal-to-background ratio (SBR) (**Figure S2B**), reduced background intensity (**Figure S2C**), and improved signal intensity tuning as measured by full width at half maximum (FWHM) (**Figure 2D**), whereas no observable improvements in staining were seen at 647 nm following quenching treatments as expected for a wavelength that has minimal autofluorescence (**Figure S2E-G**). Representative intensity histograms comparing the PBS control with each quenching condition are provided in **(Figure S2H).** Overall, these quenching methods enable fine details of IBA1^+^ microglia to be clearly seen in formic acid-treated brains (**Figure S2A)**. While LED treatments were slightly superior at improving staining quality, SBB treatment was much quicker and reproducible (20 min vs 24 hour) hence it was selected for immunofluorescent staining of molecular markers visualised at green wavelengths.We next set out to identify those markers and brain regions that most strongly associate with disease progression and to identify the most sensitive indicators of acceleration or slowing of neurodegenerative processes that could be used in future studies.

**Figure 2.**
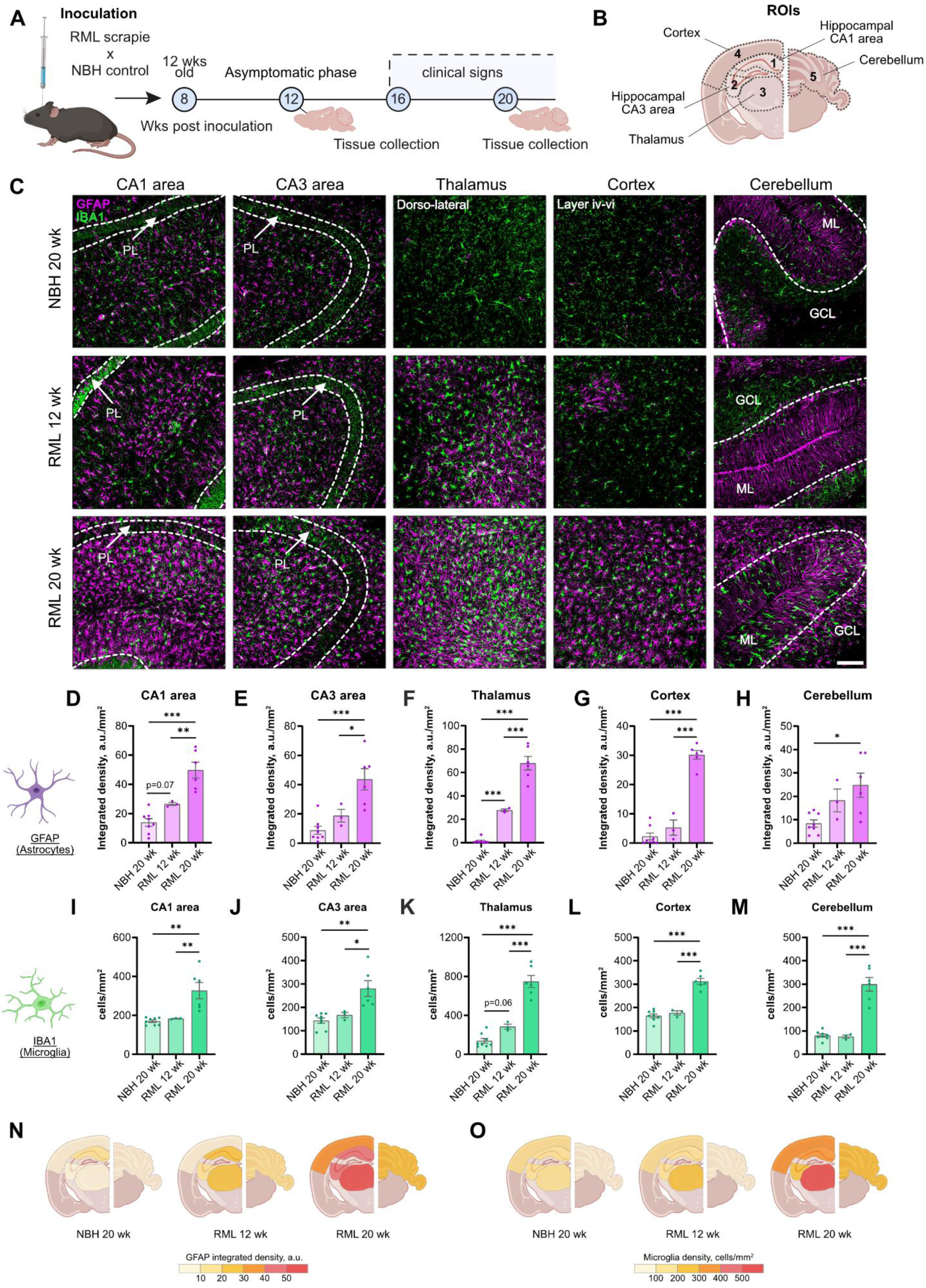
Brain-wide reactive gliosis during RML prion disease. A,B) Brains were collected at 12-wk (asymptomatic phase) and 20-wk (clinical phase) post inoculation (A), to analyse regions of interest (ROIs) relevant to neurodegeneration (B). **C)** Representative images of astrocytic GFAP (magenta) and microglial IBA1 (green) immunofluorescent co-staining in five brain ROIs. **D-H)** GFAP⁺ integrated density per 1 mm² in the CA1 area (D), CA3 area (E), thalamus (F), cortex (G), and cerebellum (H). **I-M)** Density of IBA1^+^ microglia per 1 mm² in the CA1 area (I), CA3 area (J), thalamus (K), cortex (L), and cerebellum (M). **N,O)** Brain maps are colour coded by quantified immunoreactivity for GFAP^+^ (N) or IBA1^+^ cell (O) density in NBH 20 wk, RML 12 wk, and RML 20 wk groups across ROIs. Abbreviations: PL, pyramidal layer; ML, molecular layer; GCL, granule cell layer. Significance is indicated as p < 0.05 (*), p < 0.01 (**), p < 0.001 (***). Data is presented as mean ± SEM. The scale bar shown represents 100 µm.

### RML scrapie triggers widespread astrocyte and microglia activation

To gain insight into RML scrapie-induced neuroinflammatory processes using refined immunohistochemical approaches in decontaminated prion samples, we stained for GFAP to quantify astrocyte reactivity, and IBA1 to quantify microglial cell number. We analysed the brains of mice at 12-wks (disease is present without behavioural signs) and 20-wks post inoculation with RML scrapie (behavioural signs are present) and compared these to control mouse brains at 20 weeks post inoculation with normal brain homogenate (NBH) (**Figure 2A)**. At these time points, we analysed key neurodegeneration-associated brain regions likely of greatest relevance to the community outside of the prion field (Forner et al. 2021). Specifically, we focused on the hippocampus, thalamus, cortex, and cerebellum in order to uncover region-specific vulnerability to the prion disease (**Figure 2 B**) (Kampmann 2024; DeArmond et al. 1997). We analysed the CA1 and CA3 subregions of the hippocampus separately due to their differential susceptibility to neuropathological interventions (Shipton et al. 2022; Einenkel and Salameh 2024).

We found clear immunoreactivity across each of these brain regions (**Figure 2C, S2I,J**). Using a fixed threshold to quantify immunoreactivity across each of these brain regions, we found a 3- to 13-fold elevation in GFAP^+^ integrated density at 20 weeks compared to NBH controls (**Figure 2D-H**). For example, we found that GFAP was almost undetectable in the thalamus of NBH-inoculated control mice, but was significantly increased in the brains of RML 12-wks mice, and much higher still by 20-wks (**Figure 2C,F**). We observed a similar pattern for IBA1^+^ cell density, with no changes detectable at 12 weeks, but a significant 2- to 5-fold increases seen across all examined brain regions at 20-wks (**Figure 2I-M**). The changes are summarised in brain maps for GFAP integrated intensity (**Figure 2N)** and for IBA1^+^ microglial cell density (**Figure 2O).**

### Prion inoculation induces regional-specific patterns of the ER stress marker p-PERK

Prion diseases including RML scrapie are characterised by the accumulation of misfolded protein, causing ER stress and triggering an unfolded protein response (UPR) through the PERK-eIF2α pathway (Moreno et al. 2012) . We therefore hypothesised that we would observe a widespread increase in ER stress, as visualised by immunostraining for phosphorylated PERK (p-PERK), across the RML scrapie-inoculated mouse brain. To study the spatial and temporal patterns of ER stress in RML scrapie-inoculated mice, we first applied a heat-induced antigen retrieval method to enhance the detection (p-PERK by immunofluorescence. Antigen retrieval robustly increased the signal intensity of p-PERK^+^ staining in the CA1 pyramidal layer of both NBH and RML animals (**Figure 3A, S3A-E**). We used this protocol to immunostain brain sections from NBH, RML 12-wk, and 20-wk mice with p-PERK (and Hoechst or NeuroTrace counterstaining), to quantify the level of ER stress in analysed brain ROIs (**Figure 3A, S3E**). We found that p-PERK was particularly elevated in the pyramidal neurons of hippocampal CA1 and CA3 in RML 20-wk mice, being ∼35 and ∼26 times higher, respectively than NBH controls. In contrast, p-PERK immunoreactivity was only modestly elevated in deep cortical neuronal layers and unchanged in the thalamus and cerebellum (**Figure 3B-F**). While we had expected a generalised increase in p-PERK immunoreactivity by 12-wks post-inoculation, compared to NBH in these regions, the results did not reach statistical significance. The data are summarised in brain maps (**Figure 3G**).

**Figure 3.**
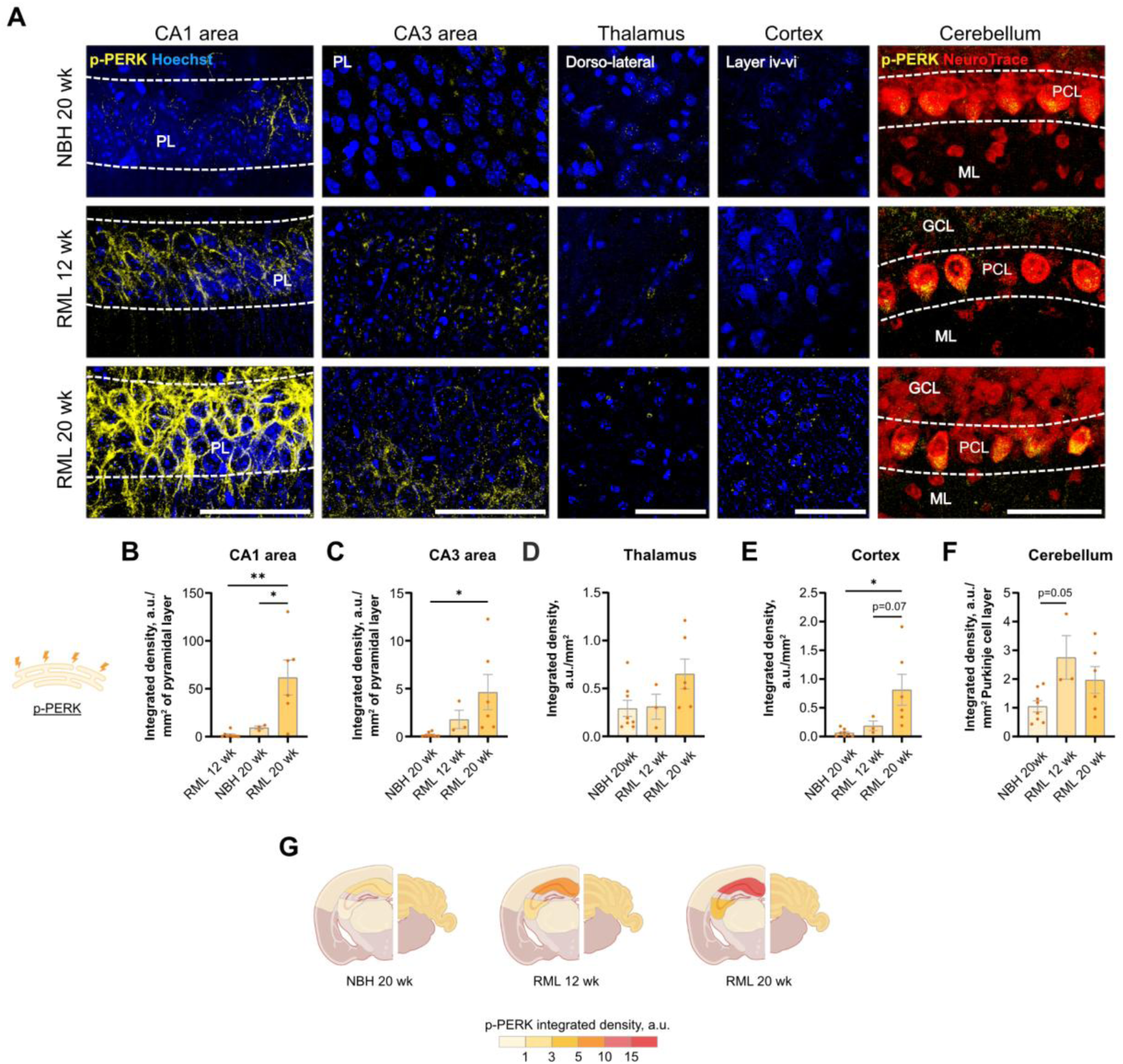
Region-specific patterns of an ER stress marker in the brains of RML scrapie-infected mice. **A)** Representative images of p-PERK immunofluorescent staining (yellow) and Hoechst (blue) or NeuroTrace 660 co-staining (red) in five brain ROIs. **B-F)** p-PERK⁺ integrated density per 1 mm² in the pyramidal layer of CA1 area (B), pyramidal layer of CA3 area (C), thalamus (D), cortex (E), and Purkinje cell layer of cerebellum (F). **G)** Brain maps are colour coded by quantified immunoreactivity for p-PERK in NBH 20 wk, RML 12 wk, and RML 20 wk groups across ROIs. Abbreviations: PL, pyramidal layer; ML, molecular layer; PCL, Purkinje cell layer; GCL, granule cell layer. Significance is indicated as p < 0.05 (*), p < 0.01 (**). Data is presented as mean ± SEM. The scale bar shown represents 50 µm.

### Lysosomal pathway dysregulation appears in late-stage prion disease

A key histopathological feature specific to prion disease is the sponge-like appearance of the brain tissue due to the accumulation of roughly spherical holes in the brain. This spongioform degeneration is thought to result from the disruption of endo-lysosomal pathways and subsequent vacuolation (Lakkaraju et al. 2021). We therefore tested for alterations in the immunoreactivity of the lysosomal marker LAMP1 as a measure of lysosomal activity and/or dysfunction in NBH, 12-wk and 20-wk mice. In contrast to p-PERK staining, we found that LAMP1 immunoreactivity was widespread in all brain regions we analysed (**Figure 4A-F, S4**). Counter-staining with Neurotrace suggested that LAMP1 was found within both neurons and non-neuronal cells, for example as seen in the hippocampal pyramidal cell layer and CA1/CA3 and cerebellar molecular layer (**Fig. 4A**). Collectively, we found that LAMP1 expression did not significantly change at the early stage of disease. However, it was strongly activated by 20-wks, particularly in the thalamus where it was almost 20 times greater compared with the NBH 20-wk group. The presence of LAMP1 is consistent with the premise of dysregulated lysosomal trafficking in RML-scrapie and coincides with an appearance of thalamic GFAP^+^ astrocytes (**Fig. 2F**). The data are summarised in brain maps (**Figure 4G**).

**Figure 4.**
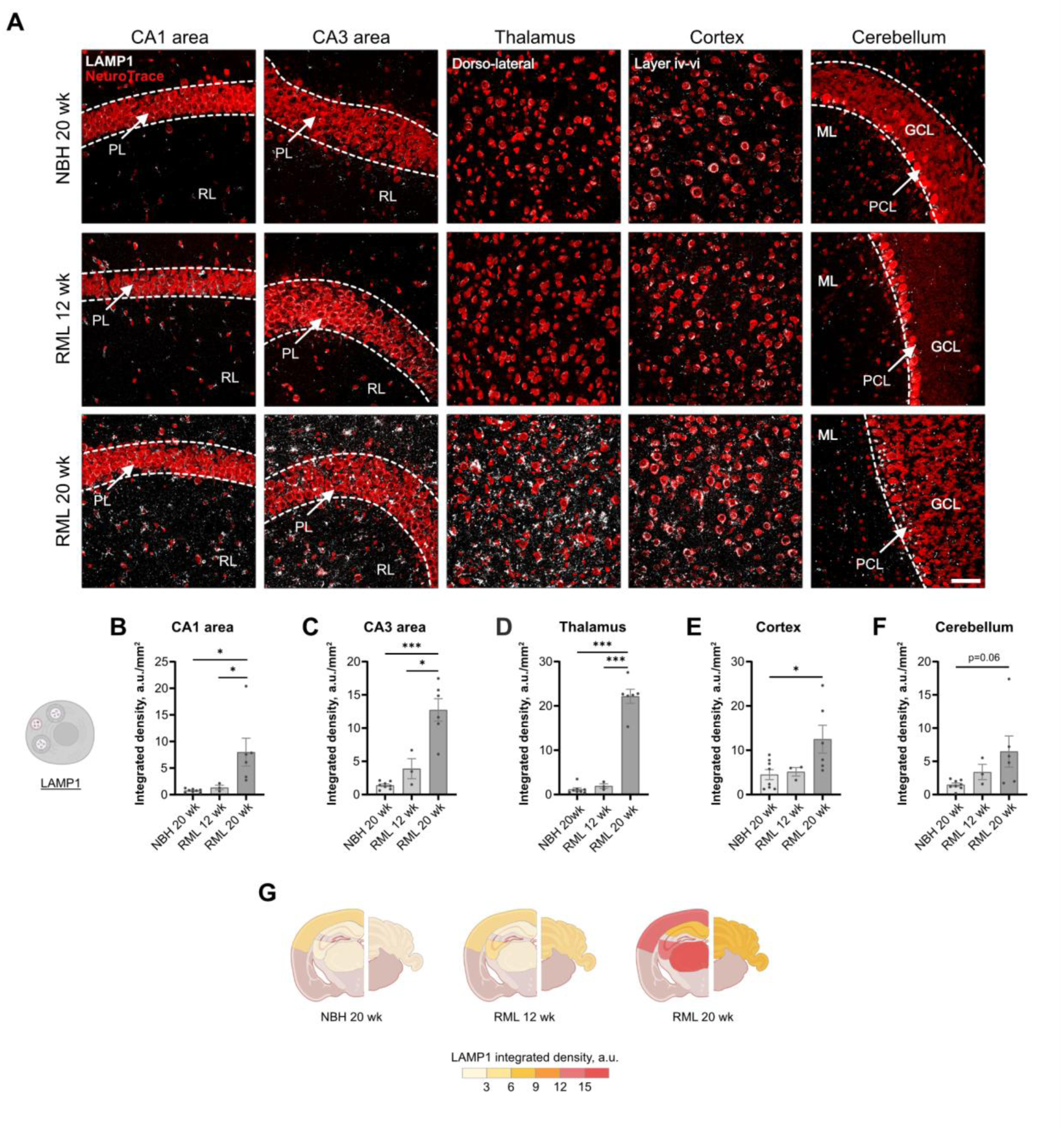
Expression of the lysosomal marker LAMP1 throughout prion disease in mice. **A)** Representative images of LAMP1 immunofluorescent staining (grey) and NeuroTrace 660 co-staining (red) in five brain ROIs. **B-F)** LAMP1⁺ integrated density per 1 mm² in the CA1 area (B), CA3 area (C), thalamus (D), cortex (E), and cerebellum (F). **G)** Brain maps are colour coded by quantified immunoreactivity for LAMP1 in NBH 20 wk, RML 12 wk, and RML 20 wk groups across ROIs. Abbreviations: PL, pyramidal layer; RL, radial layer; ML, molecular layer; PCL, Purkinje cell layer; GCL, granule cell layer. Significance is indicated as p < 0.05 (*), p < 0.00 (***). Data is presented as mean ± SEM. The scale bar shown represents 50 µm.

### Neuronal loss is observed in all brain regions irrespective of p-PERK and LAMP1 expression

The accumulation of misfolded protein in RML mice drives p-PERK, LAMP1, and GFAP expression as well as broad microglial activity consistent with induced ER stress, perturbed lysosomal function, and inflammation. We wondered how the presence of these markers would influence the process of neuronal death, since widespread neuronal loss is a key factor in the development of clinical behavioural signs of prion disease. We therefore quantified the number of NeuN immunoreactive neurons present in the hippocampal pyramidal layer, dorso-lateral thalamus, somatosensory cortical layers iv-vi, and the cerebellar granule cell layer of NBH, 12-wks, and 20-wks mice (**Figure 5A).** This analysis showed a clear reduction in neuronal density in all brain regions of interest in RML 20-wk mice, with the highest relative losses observed in the thalamus and cerebellum with approximately 50% and 25% fewer stained neurons, respectively. Surprisingly, this reduction occurred despite minimal to no detection of p-PERK in either region (thalamus or cerebellum) or of LAMP1 immunoreactivity in the cerebellum (**Figure 5B-F**). As expected, 20-wks mice had greater neuronal loss than at 12-wks in hippocampal CA1, CA3, thalamus and cerebellum but not cortex (**Figure 5B,C,F**). Overall, large regional variation in neuronal loss may suggest region-specific neuronal vulnerability in RML scrapie mice (**Figure 5G**). This variation appears to correlate with the differential expression of disease biomarkers including GFAP and LAMP1.

**Figure 5.**
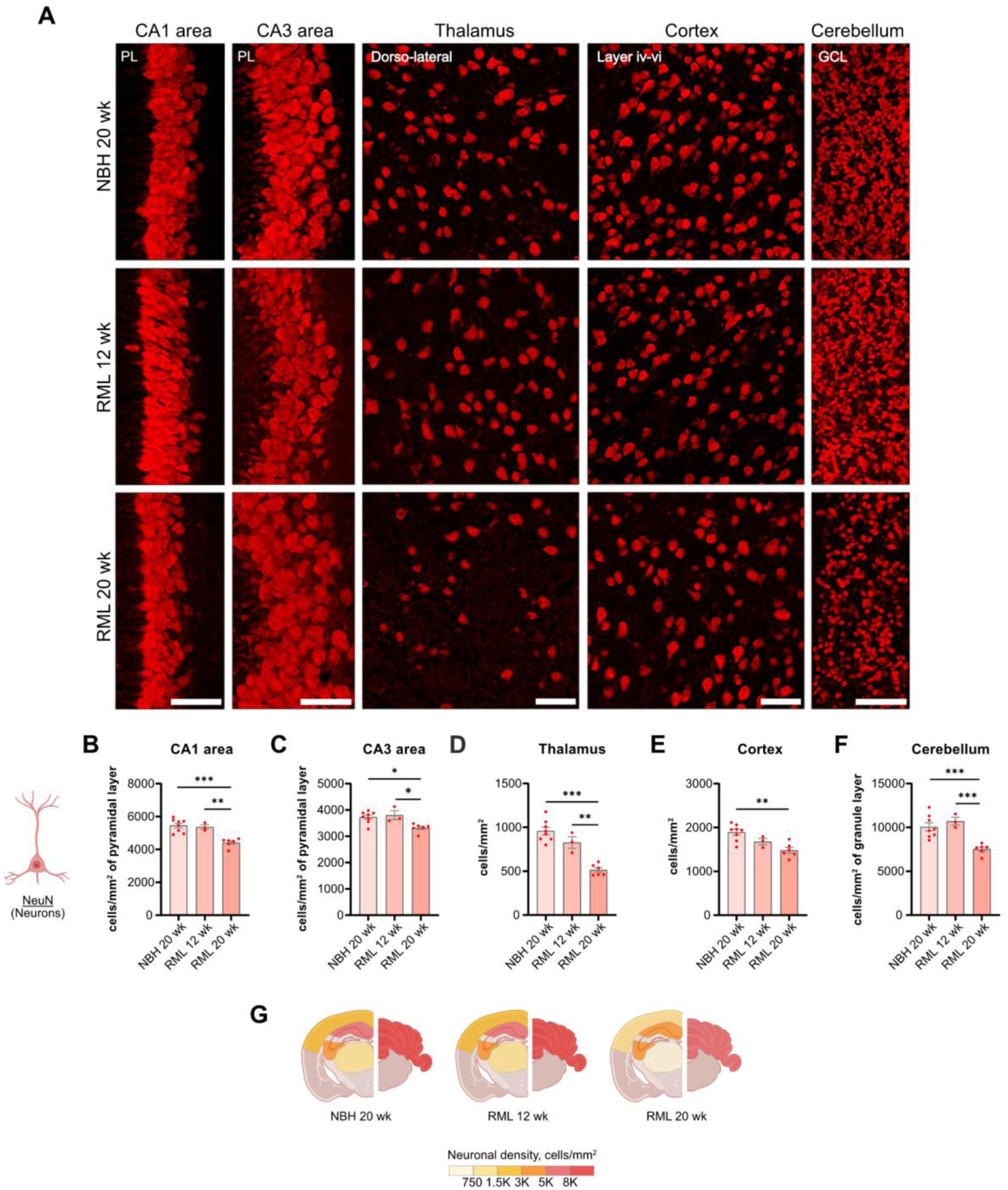
Mapping neuronal loss across brain regions in prion disease. **A)** Representative images of NeuN immunofluorescent staining in five brain ROIs. **B-F)** Density of NeuN^+^ neurons per 1 mm² in the pyramidal layer of CA1 area (B), pyramidal layer of CA3 area (C), dorso-lateral thalamus (D), somatosensory cortex (E), and granule cell layer of cerebellum (F). **G)** Brain maps are colour coded by quantified NeuN^+^ neuronal density in NBH 20 wk, RML 12 wk, and RML 20 wk groups across ROIs. Abbreviations: PL, pyramidal layer; GCL, granule cell layer. Significance is indicated as p < 0.05 (*), p < 0.01 (**), p < 0.001 (***). Data is presented as mean ± SEM. The scale bar shown represents 50 µm.

### Automated quantification reveales regional differences in spongioform degeneration

Given the widespread increase in LAMP1 expression and regional selectivity in neuronal loss we observed, we wondered if these measures influenced the appearance of spongiosis. To quantify spongioform degeneration, we embedded and cut 5-μm coronal sections from paraffin-embedded blocks generated from the same brains used for immunohistochemistry and stained these thin sections with H&E to reveal cellular structures and developing vacuolation (**Figure 6A**). Since the degree of spongiosis is typically assessed using a ‘clinical severity scale’ that has limited power to detect subtle quantitative differences between brain regions or treatment conditions (Hartmann et al. 2019), we first developed an automated spongiosis quantification pipeline. Specifically, we processed images of H&E-stained sections from both NBH and RML 20-wk mouse brains (**Figure 6B**) and manually annotated features corresponding to either true spongiosis events, blood vessels, dendrites and/or histology artefacts. We then trained a supervised machine learning model on these data using a random forest method (ilastik software) to automatically segment true positive vacuoles in a test dataset (**Figure 6C)**. And after model refinement, only a negligible number of false positive events were detected in control tissues. We then applied this model to a full dataset of 291 images for CA1 and 187 for thalamus to quantify vacuole number and size automatically (**Figure 6D** and **E).** Images were taken from both the CA1 area of the hippocampus and the thalamus since these regions showed significant but distinct molecular patterns of prion histopathology (**Figure 6F-I)**.

**Figure 6.**
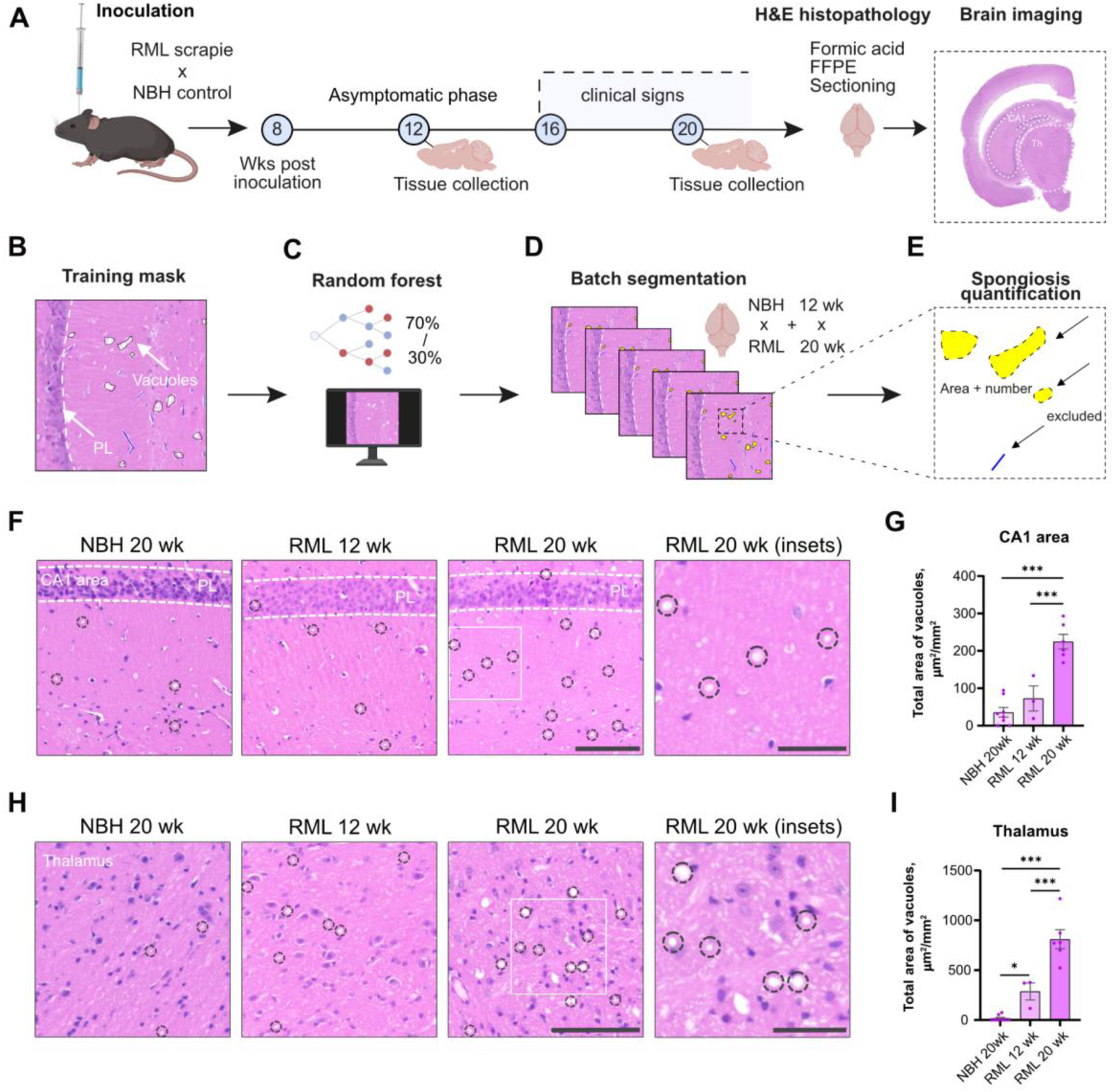
Automated approach for evaluating spongioform degeneration in RML scrapie-infected brains. **A)** The workflow for assessing spongiform degeneration included collecting brains from mice at 12 wk and 20 wk post-prion disease modelling, followed by H&E staining of brain sections and whole-brain imaging using a slide scanner. **B-E)** Sample images manually annotated with NBH and RML vacuole mask (B) used to train a random forest model for automated annotation (C) followed by batch annotation of all NBH and RML images (D)and quantification of segmented vacuoles (E). **F)** Representative images of spongiosis in the hippocampal CA1 area. **G)** Total area of brain vacuolation per 1 mm^2^ in the CA1 area. **H)** Representative images of spongiosis in the thalamus. **I)** Total area of brain vacuolation per 1 mm^2^ in the thalamus. Abbreviations: PL, pyramidal layer. Significance is indicated as p < 0.05 (*), p < 0.001 (***). Data is presented as mean ± SEM. The scale bar shown represents 100 µm and 40 µm (RML 20 wk insets).

We found that the total area of vacuolation per mm^2^ of brain tissue in the CA1 area showed a non-significant trend to increase at 12-wks post-inoculation relative to NBH, and was fourfold greater than control by 20-wks (**Figure 6G**). Remarkably, spongiosis was clearly evident in the thalamus by 12-wks to an extent equivalent to the 20-wk timepoint in CA1 (∼200 µm^2^/mm^2^), suggesting that it is either significantly more sensitive to lysosomal disruption or that scrapie misfolding or deposition is elevated in the thalamus in this model. Thalamic spongiosis was threefold greater in 20 wks (750 µm^2^/mm^2^) than either 20-wk CA1 spongiosis or 12-week thalamic spongiosis (**Figure 6I).** The extent of spongiosis in the thalamus appeared to correlate with both the loss of NeuN immunoreactivity and the progressive increases in LAMP1, GFAP, and IBA1 immunoreactivity, but not with p-PERK immunoreactivity that was prominent in CA1. Overall, this pipeline to quantify spongiosis, in combination with previously described immunohistochemical markers, suggests regional differences in vulnerability and distinct temporal patterns of neurodegeneration-associated histopathological features of prion disease.

### Region-specific vulnerability and a role for astrocytic-LAMP1 in prion disease progression

To facilitate the side-by-side comparison of histopathological markers of neurodegenerative disease in the RML scrapie-inoculated mouse brain, we normalised immunohistochemical data to the values observed in NBH controls for each analysed feature at both 12-wks and 20-wks post-inoculation (**Figure 7A).** This analysis revealed particularly large and highly significant changes in thalamic GFAP expression at 12-wks, and more widespread changes evident by 20-wks including LAMP1 in the thalamus and p-PERK in CA1 and CA3 areas. These findings are consistent with these timepoints representing separate phases of disease, and suggest that specific region and marker combinations are more sensitive indicators of early disease-related changes. Global analysis of the neuronal marker NeuN confirmed the absence of significant neuronal loss at 12-wks but significant and often dramatic increases by 20-wks, with the thalamus being most strongly affected (**Figure 7B**).

**Figure 7.**
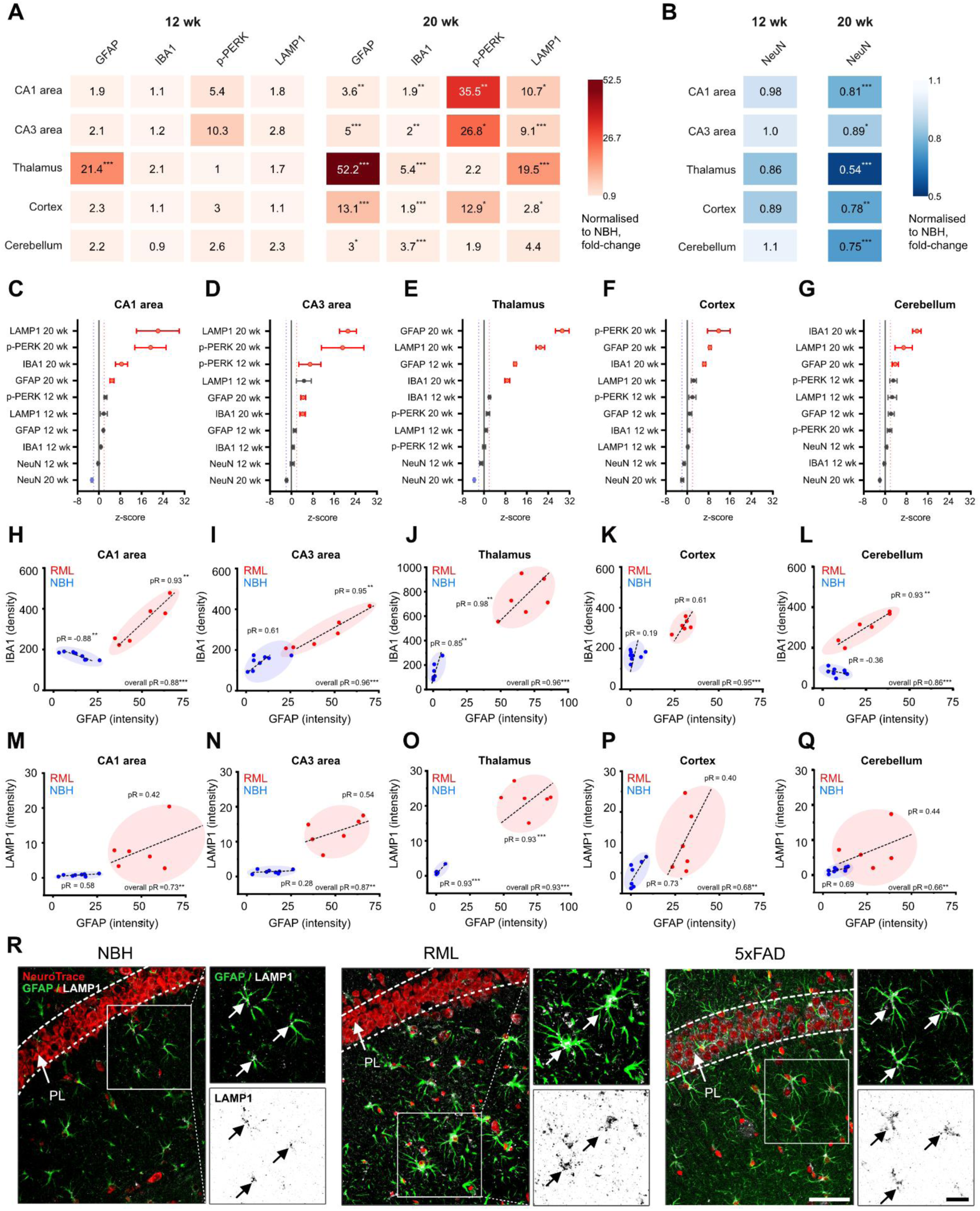
Integration of histopathological markers enables regional profiling of prion disease in the brain. A-B) Heat maps of immunohistochemical markers that were upregulated (A) and downregulated (B) in scrapie-induced neurodegeneration. **C-G)** Plots of ranked z-scores relative to NBH for all markers and time points analysed in the hippocampal CA1 area (C), hippocampal CA3 area (D), thalamus (E), cortex (F) and cerebellum (G). **H-L)** Correlative plots between GFAP intensity and IBA1^+^ microglia density in the hippocampal CA1 area (H), hippocampal CA3 area (I), thalamus (J), cortex (K) and cerebellum (L) of NBH 20 wk and RML 20 wk mice. **M-Q)** Correlative plots between GFAP and LAMP1 intensity in the hippocampal CA1 area (M), hippocampal CA3 area (N), thalamus (O), cortex (P) and cerebellum (Q) of NBH 20 wk and RML 20 wk mice. **R)** Illustrative images showing GFAP and LAMP1 co-localisation in the radial layer of the hippocampal CA1 area in NBH, RML, and 5xFAD mice. Abbreviations: PL, pyramidal layer. Significance is indicated as p < 0.05 (*), p < 0.01 (**), p < 0.001 (***). Data is presented as mean ± SEM. The scale bars shown represent 50 µm (hippocampal CA1 area) and 20 µm (GFAP/LAMP1 magnified crops).

We therefore wondered whether some regions of the brain had greater apparent responses to prion inoculation, and whether different markers were optimal for staging progression in different regions. To this end, we took the ranked z-score relative to NBH as a measure of signal-to-noise for all markers, timepoints, and regions of interest we analysed (**Figure 7C-G**). We found that LAMP1 had the highest score in CA1 and CA3 (**Figure 7C,D**), and the second highest in cortex and cerebellum (**Figure 7F,G**) while p-PERK ranked first or second in CA1, CA3 and the cortex (**Figure 7C,D,F**) but surprisingly was not significant in the thalamus where signals from GFAP and LAMP1 dominate (**Figure 7E).** As seen in spongiosis data, the thalamus (**Fig. 6I**) was also most greatly associated with accumulating vacuoles at early and late timepoints.

In prion models, the p-PERK-eIF2α ER stress pathway is well-known to induce translational arrest and apoptosis under specific conditions (Albert-Gasco et al. 2024; Mays and Soto 2016). Here, we observed strong thalamic neuronal loss in response to prion inoculation, accompanied by spongiosis, without observing p-PERK immunoreactivity that was present in other brain regions. The thalamus was also unique in the early expression of GFAP in astrocytes. Thus we sought to better understand this relationship by looking at region-specific correlations with GFAP focusing on IBA1 as a marker of reactive microgliosis as well as LAMP1 as a marker of lysosomal dysfunction. As expected, in NBH-inoculated animals, IBA1 and GFAP both tend towards the zero intercept in all regions (**Figure 7H-L**). This relationship changes significantly when disease is present when IBA1 density correlates significantly (pR > 0.93), with increased astrocytes immunoreactivity in all analysed brain regions (**Figure 7H-J,L**) other than the cortex (**Figure 7K**). When we considered the overall correlations, including both groups, we observed significant correlations between LAMP1 and GFAP in all brain regions (**Figure 7 M-Q).** Most interestingly, in the thalamus we found that LAMP1 was positively correlated (pR > 0.93) to astrocytic GFAP in RML-inoculated mice but not NBH mice alone (**Fig. 7O)**.

As we observed a particularly strong relationship between LAMP1 and GFAP expression in the thalamus (**Fig. 7O)**, we wondered whether these markers might be linked biologically. We therefore co-stained for GFAP and LAMP1 in brains of NBH 20-wk, RML 20-wk, 5xFAD mice (familial AD) to test the hypothesis that these two markers co-localise across multiple proteinopathy models. Upon counterstaining with NeuroTrace to label neurons and high-magnification confocal imaging of the hippocampus, we found clear evidence for co-localisation of LAMP1 within GFAP+ cells with astrocytic morphology (**Figure 7R**). Furthermore, astrocytic LAMP1 immunoreactivity appeared to be higher in both RML 20-wk and 5xFAD mice. Together, these findings suggest that LAMP1 accumulation in astrocytes may contribute to prion disease pathology and may have a more general role in neurodegenerative proteinopathies including AD.

In summary, we have mapped the progression of prion disease markers including GFAP, IBA1, p-PERK, LAMP1, spongiosis formation, and neuronal loss across multiple brain regions at both early and late disease timepoints. Overall, we found that pyramidal cells of the hippocampal CA1 and CA3 areas appear particularly vulnerable to p-PERK accumulation while the thalamus lacks obvious p-PERK reactivity but displays significant glial activation, lysosomal disruption as suggested by increased spongiosis and LAMP1 immunoreactivity, and neurodegeneration.

## DISCUSSION

In this study, we report the refinement of the well-established and rapid RML scrapie model of neurodegeneration by establishing protocols for paired tissue collection, decontamination, and high-quality brain immunohistochemistry at just 20 wks post inoculation in WT mice. Using this model, we identified region-specific vulnerabilities and key markers that best stage RML scrapie-induced neurodegenerative disease progression. We found that the markers GFAP and LAMP1 are associated with the development of key pathological outcomes including neuronal loss and spongiosis, and are particularly apparent in the hippocampus and thalamus. We also provide evidence for a shift from predominantly neuronal LAMP1 to astrocytic LAMP1, implicating the astrocytic-lysosomal pathway in neurodegenerative processes.

The protocols and methods we describe enable simpler and more robust molecular staging of disease as an alternative to traditional survival protocols (Harding et al. 2024; Nuvolone et al. 2015; Callender et al. 2020). Carrying out analysis at relatively short time points during gRML scrapie progression also facilitates faster experimental iteration and reduced animal suffering side effects such as bladder enlargement that complicate studies with female mice (Guenther et al. 2001). Since the model requires no significant breeding or genetic modification, it enables well-powered preclinical studies and is suitable for paired-sample biochemistry and the integration of multi-omics data. We hope our characterisation of this protocol will facilitate the wider adoption of an underutilised and powerful model for studying neurodegeneration and developing new therapies.

### Key findings and future directions

We have identified region-specific patterns of RML scrapie-associated markers that differentially correlate to neuronal loss. These findings suggest that while neurodegeneration is widespread, different mechanisms may be at play in different areas of the brain. Our integrated analysis of quantitative histological data suggests that the thalamus and hippocampal CA1 and CA3 areas have the highest pathological burden. Consistently with our result, multiple studies have reported that these regions also show the highest level of PrP^Sc^ accumulation following intracerebral RML prion inoculation in mice (DeArmond et al. 1997; Moda et al. 2015; Foliaki et al. 2024).

Neuronal loss was notable in all regions by 20-wks, but was particularly high (almost 50% loss) in the thalamus and coinciding with significantly increased LAMP1 immunoreactivity and spongioform degeneration. Impaired lysosomal trafficking is a plausible explanation for vacuolisation and neuronal loss and can arise from a well characterised pathway involving PIKfyve deacylation, driven by ER stress (Lakkaraju et al. 2021). However, we were unable to detect significantly elevated p-PERK in the thalamus as evidence in support of this pathway. Furthermore, spongiosis was observed at 12-wk post-inoculation, alongside GFAP^+^ astrocytes, but prior to LAMP1 detection, leaving the causative role of LAMP1 in vacuole formation in doubt. We note that, in our hands, p-PERK immunoreactivity was predominantly neuronal but we do not rule out UPR activation in the non-neuronal populations that has been reported (Smith et al. 2020). This work also supports the role of autophagy pathways in prion disease.

In the healthy brain, LAMP1 expression was observed in CA1 and CA3 pyramidal neurons, cortical neurons and Purkinje neurons. However, during RML scrapie, LAMP1 expression was clearly detectable in GFAP^+^ cells with astrocytic morphology in the CA1 area, thalamus, and granule cell layer. This expression coincided with a dramatic increase in GFAP^+^ immunoreactivity across the brain (and the restricted appearance of p-PERK in neurons) and was particularly notable in the thalamus. The role of GFAP^+^-LAMP1^+^ astrocytes is unclear but is consistent with similar patterns of expression reported in 5xFAD mice (Gowrishankar et al. 2015). Similarly, a substantial upregulation of LAMP1 in hippocampal astrocytes in the APP/PS1 mouse model of Alzheimer’s Disease has been reported, as well as in primary astrocyte cultures challenged with Aβ oligomers ((Li et al. 2024). When hippocampal CA1 astrocytes show high levels of p-PERK and p-eIF2α (not observed directly in our study) they have been shown to express LAMP1 and show LC3-II–associated vacuolization (Kim et al. 2017). Similarly, LAMP1^+^ astrocytic lysosomes have been shown to co-locolised with engulfed synapses *in vivo* (Li et al. 2024), and in the spinal cord, GFAP^+^-LAMP1^+^ astrocytes co-express the death receptor TRAIL (TNF-related apoptosis-inducing ligand), and can induce apoptosis (Sanmarco et al. 2021). Lastly, in a comparable prion model, disruption of the astrocyte secretome by the PERK-eIF2α pathway led to synaptic loss and neuronal death (Smith et al. 2020). We suggest there may be an important transition from neuronal LAMP1 to astrocytic LAMP1 that may contribute to neurodegenerative processes, and this mechanism warrants further investigation.

In contrast to the hippocampus and thalamus, the cerebellum did not show significant upregulation of either LAMP1 nor p-PERK despite a substantial reduction (∼25%) in neuronal number in the granule cell layer. This finding is consistent with previous reports of cell-type specific vulnerability in both prion-infected and AD-related cell types driven by differential expression of key genes such as Rbm3 (Slota et al. 2024). Notably, *Rbm3* was previously identified as strongly neuroprotective in prion-inoculated mice, suggesting its expression be important for region-specific differences in the neuronal loss we observed (Peretti et al. 2015; Preußner et al. 2023). Lastly, we noted widespread and often highly correlated IBA1 and GFAP immunoreactivity across the brain of RML mice, indicative of a widespread inflammatory response. This expression pattern contrasted to p-PERK and LAMP1 that had clear spatial patterns of expression (e.g. pyramidal or purkinje cells) within each region.

### Best practice in the safe handling of scrapie-infected samples

Models of prion disease understandably raise concern due to a history of bovine spongiform encephalopathy (BSE) (Houston and Andréoletti 2019), and the rare but catastrophic laboratory transmission of variant Creutzfeldt–Jakob disease (CJD) to humans (Brandel et al. 2020). In contrast, evidence shows that RML scrapie provides the benefits of an aggressive prion model with minimal risk. Specifically, there is no evidence of transmissibility potential of scrapie isolates to humans despite a long history of ovine meat in the food chain (EFSA Panel on Biological Hazards (BIOHAZ Panel) 2015; Cann 2016). Transmission of scrapie to transgenic mice expressing humanised prion protein (TgHu) does not demonstrate clinical signs on primary inoculation requiring serial passaging (brain homogenate from one mouse to the next) to produce the infectious prion strain (Wadsworth et al. 2013; R. Wilson et al. 2012; Houston and Andréoletti 2019). To date, transmission of scrapie to a single cynomolgus macaque has been achieved only by direct inoculation to the brain with a very high dose of infected brain tissue (25 mg), an entirely artificial scenario in the context of laboratory safety (Comoy et al. 2015). These studies were all performed without decontamination. We argue that when used in appropriate Category 2 facilities (Advisory Committee on Dangerous Pathogens (ADCP) 2015) and combined with well validated decontamination procedures using formic acid (Taylor et al. 1997; Brown, Wolff, and Gajdusek 1990; Tateishi, Tashima, and Kitamoto 1991; Dong et al. 2021) that are also used for Category 3 human CJD samples, RML scrapie mice and their tissues are a a safe and powerful model to study neurodegeneration.

### Challenges in modelling neurodegenerative diseases

The RML scrapie model and the methods we describe l focuses on the dissection of neurodegenerative pathways downstream of PrP^sc^ accumulation. This model can be used to study the response of neurons and other cell types to misfolded protein, but it is not a model of AD or other human dementia. Historically, familial inherited forms of dementia with well-characterised genetics have been leveraged to produce mouse models that recapitulate specific features of sporadic dementia. For example, mutations in PSEN1/2, APP that lead to familial Alzheimer’s Disease (FAD) (Andrade-Guerrero et al. 2023), are overexpressed in canonical AD mouse lines such as the *App*^NL-G-F^. Alternative disease models include the more recent tau triple mutant FTD mouse line (e.g MAPT^P301S;Int10+3;S320F^). The introduction of these mutations allows key clinical and pathological signs seen in patients to be reproduced in mice. Similar research has been conducted on humanised mice carrying genetic risk factors for sporadic Late Onset Alzheimer’s Disease (LOAD) such as APOE ε4 (Robinson et al. 2018). These models have resulted in significant mechanistic success alongside the well described challenges in translating treatments over many years into patients (Qian, Li, and Ye 2024). While each of these models recapitulates aspects of neurodegenerative disease, linking the diversity of mechanisms resulting from each model to human disease is challenging. The RML scrapie model is, in our view, undervalued in this context precisely because the mechanism of protein accumulation and misfolding is much simpler and better understood, can be carried out in WT mice eliminating the need for complex breeding paradigms, and progresses relatively rapidly producing stereotyped phenotypes.

We hope that the methods and deeper histopathological characterisation we provide in this study will promote the use of this model for mechanistic discovery and scalable pre-clinical drug testing for neurodegenerative disease. The fundamental processes behind neurodegeneration are not well understood but, we believe, may be common to all neurodegenerative conditions (D. M. Wilson 3rd et al. 2023). Thus studying these processes efficiently, in interpretable models, may provide key insights into urgently needed mechanisms therapies needed to stop neurons from dying.

## ACKNOWLEDGEMENTS

We thank Giovanna Mallucci for providing RML and NBH homogenate, the Histopathology core facility at the Institute of Metabolic Science (IMS) for paraffin embedding of brain samples, and the Tissue and Cell Imaging core facility at the IMS for assistance with imaging. The TCI is funded by the UK Medical Research Council (MRC) Metabolic Disease Unit (MRC_MC_UU_00014/5) and a Wellcome Trust Major Award (208363/Z/17/Z), FTM is a New York Stem Cell Foundation -Robertson Investigator [NYSCF-R-156] and was supported by the Wellcome Trust (WT) and Royal Society [211221/Z/18/Z]. Costs associated with this study were supported by these grants and an award from the Chan Zuckerberg Initiative [CZI NDCN 191942, 10.37921/429861umrcjh]. ECH was also supported by the Isaac Newton Trust, and DS was supported by the Ukrainian Academic Support Scheme of the University of Cambridge, and the Rowan Williams Scholarship of the Cambridge Trust. For the purpose of open access, the authors have applied a CC-BY public copyright license to any Author Accepted Manuscript version arising from this submission.

## AUTHOR CONTRIBUTIONS

**DS, ECH** and **FTM** designed experiments and drafted the manuscript with input from all authors. **DS, IAK, H-JCC** and **ECH** analysed data and produced visualisations and performed experiments. **FTM** acquired funding and assisted in data interpretation and worked with **ECH** to conceive of and manage the project and personnel.

## DATA AND CODE AVAILABILITY

All original code is deposited on Github and archived by Zenodo. The DOI is publicly available in the key resource table.

## MATERIALS AND METHODS

### Mouse husbandry

Animal work was performed under a Project Licence administered by the UK Home Office and approved by the University of Cambridge Animal Welfare and Ethics Review Board (AWERB) adhering to 3Rs principals and ARRIVE guidelines. C57BL/6J male mice used in this study were purchased from Charles River Laboratories (Saffron Walden, UK) at four weeks of age, fed a standard chow diet on arrival and housed at 3 mice per individually ventilated cage. No procedure was performed for at least seven days after arrival to enable their acclimatisation to a new facility. The animal facility’s temperature was controlled at 22 ± 2°C with a standard 12-hour light/dark cycle. Water and food were available ad libitum.

### Inoculation and treatment group randomisation

Mice were randomly allocated for unilateral inoculation in the right parietal cortex with 30 µL of a 1% w/v with either Rocky Mountain Laboratory (RML) prion (n=9) or Normal Brain Homogenate (NBH; n=8) between the ages of 4 and 6 weeks, while under isoflurane anaesthesia as previously reported (Halliday et al. 2017) monitored until recovery, and then returned to their home cage. The homogenate solution was a gift from the laboratory of Prof. Giovanna Mallucci. Mice were fed standard chow diet until 7 weeks old, then switched to a balanced control diet (10% fat diet D12450Ji, Research Diets). Mice were weighed at least once weekly.

### Terminal tissue and blood collection

At 12-wk and 20-wk post inoculation, mice were culled by the following non-recovery procedure. Under deep isoflurane anaesthesia, blood was collected by cardiac puncture using a heparin-coated (375095, Sigma Aldrich) syringe with a 23G blunt-end needle, followed by cervical dislocation. Blood was collected in tubes containing lithium heparin (450537, Greiner Bio-One MiniCollect™) and plasma was separated by 10-min centrifugation at 800 x g. Organs were dissected, transferred to cryovials, and snap-frozen on dry ice before storage at −70°C. Brains were divided along the sagittal midline, with the right hemisphere micro-dissected into different brain regions and snap frozen as above. The left hemisphere was reserved for histology by overnight fixation in 4% PFA.

### Brain decontamination

Samples were handled in accordance with guidelines from the UK Advisory Committee on Dangerous Pathogens (ADCP) and UK Health and Safety Executive. Brain hemispheres to be used for histology were washed in PBS following PFA fixation and decontaminated in 10 ml of >95% formic acid for 1 h. Following formic acid treatment, brains were subjected to a second fixation in 4% PFA overnight, washed in PBS and stored in a PBS solution containing 30% sucrose and 0.02% sodium azide at +4°C.

### Tissue sectioning

Hemispheres were cut using a Leica VT1000S vibratome (Leica Biosystems, Germany). For immunohistochemistry, 25–μm–thick coronal brain sections were cut between approximately stereotaxic coordinates 1.55 and 2.48 mm (containing the hippocampus, thalamus, and cortex) and between 6.55 and 7.65 mm (containing the cerebellum) posterior to bregma (Lein et al. 2007). Free-floating brain sections were stored in 0.05% sodium azide in PBS at +4 °C or transferred to a cryoprotectant solution (30% glycerol and 30% ethylene glycol in 0.02 M PBS) and stored at −20 °C for long-term preservation. For H&E staining, 500–600 μm–thick sections were obtained from the same hemispheres using a vibratome, embedded in paraffin, and sectioned into 5–μm slices on a Leica RM2135 microtome (Leica Biosystems, Germany). After sectioning, histology equipment was decontaminated with 2 M sodium hydroxide for 1 h and then cleaned with distilled water.

### Immunohistochemistry

Free-floating brain sections were washed in PBS prior to immunofluorescent staining. Sections designated for p-PERK staining underwent antigen retrieval in sodium citrate buffer (pH 6.0) at +90 °C for 15 min in a water bath (Grant Instruments, UK), while sections for GFAP/IBA1 staining were treated with 0.3% Sudan Black B (Alfa Aesar, USA) in 70% ethanol for 20 min to reduce background autofluorescence and improve signal-to-background ratio (Oliveira et al. 2010). Brain sections were then incubated in a blocking solution containing 5% normal donkey serum (NDS; Jackson ImmunoResearch Labs, USA), 1% bovine serum albumin (BSA; Miltenyi Biotec Ltd, UK), and 0.3% Triton X–100 (Sigma-Aldrich, USA) in PBS for 1 h at room temperature. Sections were subsequently incubated with primary antibodies diluted in blocking solution (chicken anti-GFAP, 1:1500, Antibodies.com, UK; rabbit anti-IBA1, 1:750, Abcam, UK; rabbit anti-phospho-PERK, 1:250, Cell Signalling Technology, USA; rabbit anti-LAMP1, 1:500, Abcam, UK; rabbit anti-NeuN, 1:1000, Millipore, USA) overnight at +4 °C. After three washes in PBS (30 min each), sections were incubated with appropriate secondary antibodies diluted in a blocking solution without Triton X-100 (goat anti-chicken IgY AF647, 1:1000, Invitrogen, USA; donkey anti-rabbit IgG AF488, 1:1000, Invitrogen, USA; donkey anti-rabbit IgG AF555, 1:1000, Invitrogen, USA) for 1.5 h at room temperature. Samples were counterstained with NeuroTrace 660 (1:100, Invitrogen, USA) or Hoechst 33342 (1:10000, Invitrogen, USA) for 20 min, transferred onto histological slides (Epredia, USA), air-dried for 30 min, and mounted using Aqua/Poly-Mount medium (Polysciences, USA).

### Confocal microscopy

Immunofluorescent images of mouse brains were obtained using a Leica SP8 confocal microscope (Leica Biosystems, Germany) at 20x (GFAP/IBA1), 40x (autofluorescence, NeuN, LAMP1), or 63x (p–PERK) objective magnification. Acquisition parameters were as follows: 16–bit depth; 1024 × 1024 px resolution; scan speed 400; line averaging 2; pinhole size 2.0 AU (20x) or 1.0 AU (40x and 63x); z–stacks of 3 optical sections with a z–step of 2 μm (20x) or 1 μm (40x and 63x). For autofluorescence measurements, images were acquired sequentially using laser lines at 405 nm (emission range 450/50), 488 nm (emission range 525/50), and 561 nm (emission range 629/31). Additional imaging settings for specific markers were as follows: GFAP (laser line 647 nm, emission range 672/18), IBA1 (laser line 488 nm, emission range 520/15), p–PERK (laser line 555 nm, emission range 575/15), NeuN and LAMP1 (laser line 555 nm, emission range 575/17).

### Image analysis

Morphometric analysis of 5-12 fluorescent images/region/animal was performed in ImageJ v.1.54p (NIH, USA) (Schneider, Rasband, and Eliceiri 2012) using custom macros. Z-stacks were converted into Z-projections using either the “Maximum Intensity” function or “Median” projection (for IBA1). For NeuN and p–PERK quantifications, manually defined regions of interest (ROIs) were generated using the NeuN channel (for NeuN quantification) or guided by Hoechst or NeuroTrace counterstaining (for p–PERK). Integrated density was used to quantify GFAP^+^, p–PERK^+^, and LAMP1^+^ signals after setting a fixed threshold. For IBA1^+^ microglial density analysis, Bandpass filtering (min radius = 3 px, max radius = 100 px) and Gaussian Blur (radius = 2.0) were performed prior to thresholding with the “Max Entropy” function. Analysis of particles was subsequently carried out to assess microglial density. For NeuN^+^ neuronal density, preprocessing involved Gaussian Blur (radius = 1.5) and background subtraction (“Subtract Background” with rolling ball radius = 50), followed by thresholding using either “Otsu” (hippocampus) or “Max Entropy” (thalamus, cortex, and cerebellum). Neurons were further segmented with the “Watershed” function before the analysis of particles.

### H&E staining

Super-frost histological slides containing 5 μm thick brain sections were deparaffinised in two changes of xylene (10 min each), followed by two changes of ethanol (5 min each). Slides were then stained with Mayer’s haematoxylin for 7 min, rinsed in running tap water for 5 min, and counterstained with 1% eosin (Pioneer Research Chemicals, UK) for 45 s. After dehydration in two changes of ethanol and one change of xylene (1 min each), sections were mounted under coverslips using DPX mounting medium (Sigma-Aldrich, Spain). Whole-brain images were acquired using a Zeiss Axioscan Z1 slide scanner (Zeiss, Germany) with a 20x objective.

### Spongiosis quantification

Segmentation of vacuoles for spongiosis was performed with ilastik software v.1.4.0 (Berg et al. 2019), using a random forest classifier trained on 11 manually-annotated images from 20-wk RML (47 vacuoles) and NBH-inoculated mice (88 non-vacuoles / background), and then validated on a holdout set of 7 randomly selected images (RML x 4, NBH x 3). Features included were pixel colour, intensity, texture, and edge information, with Gaussian smoothing (σ=10 pixels) applied. The classifier was re-trained iteratively as additional labelled regions were added until stable classification performance was achieved. The model was cross-validated on additional annotated images to ensure concordance with manual counts. Batch-processed outputs were analysed using a custom FIJI (ImageJ 1.54p) macro to extract vacuole size information in each image, with a minimum size filter of 5 μm² per event.

### Statistical analysis

Data was analysed in either GraphPad Prism v.8.0.2, Origin Pro 2024 or in R using custom scripts. Averaged values from 5-12 images per animal were used as biological replicates for statistical assay (n=8 in the NBH 20 wk, n=3 in the RML 12 wk, n=6 in the RML 20 wk). Analysis of one factor was performed by one-way ANOVA. The Benjamini-Hochberg FDR correction was used to control for multiple comparisons. Differences between groups were considered statistically significant at p < 0.05.

### Key resources table

| REAGENT or RESOURCE | SOURCE | IDENTIFIER |
| --- | --- | --- |
| Antibodies |  |  |
| Chicken polyclonal anti-GFAP | Antibodies.com | Cat# A85307, RRID: AB_2748894 |
| Rabbit monoclonal anti-IBA1 | Abcam | Cat# ab178847, RRID: AB_2832244 |
| Rabbit monoclonal anti-phospho-PERK | Cell Signaling Technology | Cat# 3179, RRID: AB_2095853 |
| Rabbit polyclonal anti-LAMP1 | Abcam | Cat# ab24170, RRID: AB_775978 |
| Rabbit polyclonal anti-Fox-3 (NeuN) | Millipore | Cat# ABN51, RRID: AB_10807151 |
| Goat polyclonal anti-Chicken IgY, Alexa Fluor™ 647 | Invitrogen | Cat# A-21449, RRID: AB_2535866 |
| Donkey polyclonal anti-Rabbit IgG, Alexa Fluor™ 488 | Invitrogen | Cat# A-21206, RRID: AB_2535792 |
| Donkey polyclonal anti-Rabbit IgG, Alexa Fluor™ 555 | Invitrogen | Cat# A-31572, RRID: AB_162543 |
| Biological samples |  |  |
| RML mouse-passaged scrapie homogenate | Prof. Giovanna Mallucci's lab | N/A |
| Mouse normal brain homogenate | Prof. Giovanna Mallucci's lab | N/A |
| Chemicals, peptides, and recombinant proteins |  |  |
| Normal donkey serum | Jackson ImmunoResearch | Cat# 017-000-121 |
| Bovine serum albumin 25% | Gibco | Cat# A10008-01 |
| Triton X-100 | Sigma-Aldrich | Cat# T8787 |
| Sudan Black B, powder | Alfa Aesar | Cat# J62268 |
| Copper (II) sulfate pentahydrate | Fisher Scientific | Cat# AC197722500 |
| Sodium citrate tribasic dihydrate | Sigma-Aldrich | Cat# S4641 |
| Isoflurane |  |  |
| Paraformaldehyde, powder | Sigma-Aldrich | Cat# P6148 |
| Sodium azide | Sigma-Aldrich | Cat# S2002 |
| Glycerol | Sigma-Aldrich | Cat# G5516 |
| Ethylene glycol | Sigma-Aldrich | Cat# 102466 |
| Sodium hydroxide pellets | Sigma-Aldrich | Cat# 567530 |
| Formic acid | Sigma-Aldrich | Cat# F0507 |
| Xylene | Fisher Scientific | Cat# X/0250/17 |
| Ethanol, absolute | Sigma-Aldrich | Cat# 32221 |
| Haematoxylin (Mayers) | Pioneer Research Chemicals | Cat# PRC/R/42 |
| Eosin (1% Aqueous) | Pioneer Research Chemicals | Cat# PRC/66/1 |
| NeuroTrace™ 640/660 Deep-Red Fluorescent Nissl Stain | Invitrogen | Cat# N21483 |
| Hoechst 33342, trihydrochloride, trihydrate | Invitrogen | Cat# H3570 |
| Aqua/Poly-Mount mounting medium | Polysciences | Cat# 18606 |
| DPX mounting medium | Sigma-Aldrich | Cat# 06522 |
| Experimental models: Organisms/strains |  |  |
| Mouse: C57BL/6J | Charles River Laboratories | RRID: IMSR_JAX:000664 |
| Software and algorithms |  |  |
| ImageJ v.1.54p | Schneider et al. | https://imagej.net/ij/ |
| Ilastik v.1.4.0 | Berg et al. | https://www.ilastik.org/ |
| GraphPad Prism v.8.0.2 | GraphPad Software | https://www.graphpad.com/ |
| Origin Pro 2024 | OriginLab | https://www.originlab.com/2024 |
| R (4.4.2), R Studio 2024.09.0 | Posit PBC | https://www.r-project.org |
| scrapie_image_analysis_pipelines | This paper | 10.5281/zenodo.16881829 |
| Other |  |  |
| Standard chow diet, D12450Ji | Research Diets | https://www.researchdiets.com/formulas/D12450Ji |
| LED Flood Light IP66, 100W | N/A | N/A |

## Supplementary Figures

**Figure S1.**
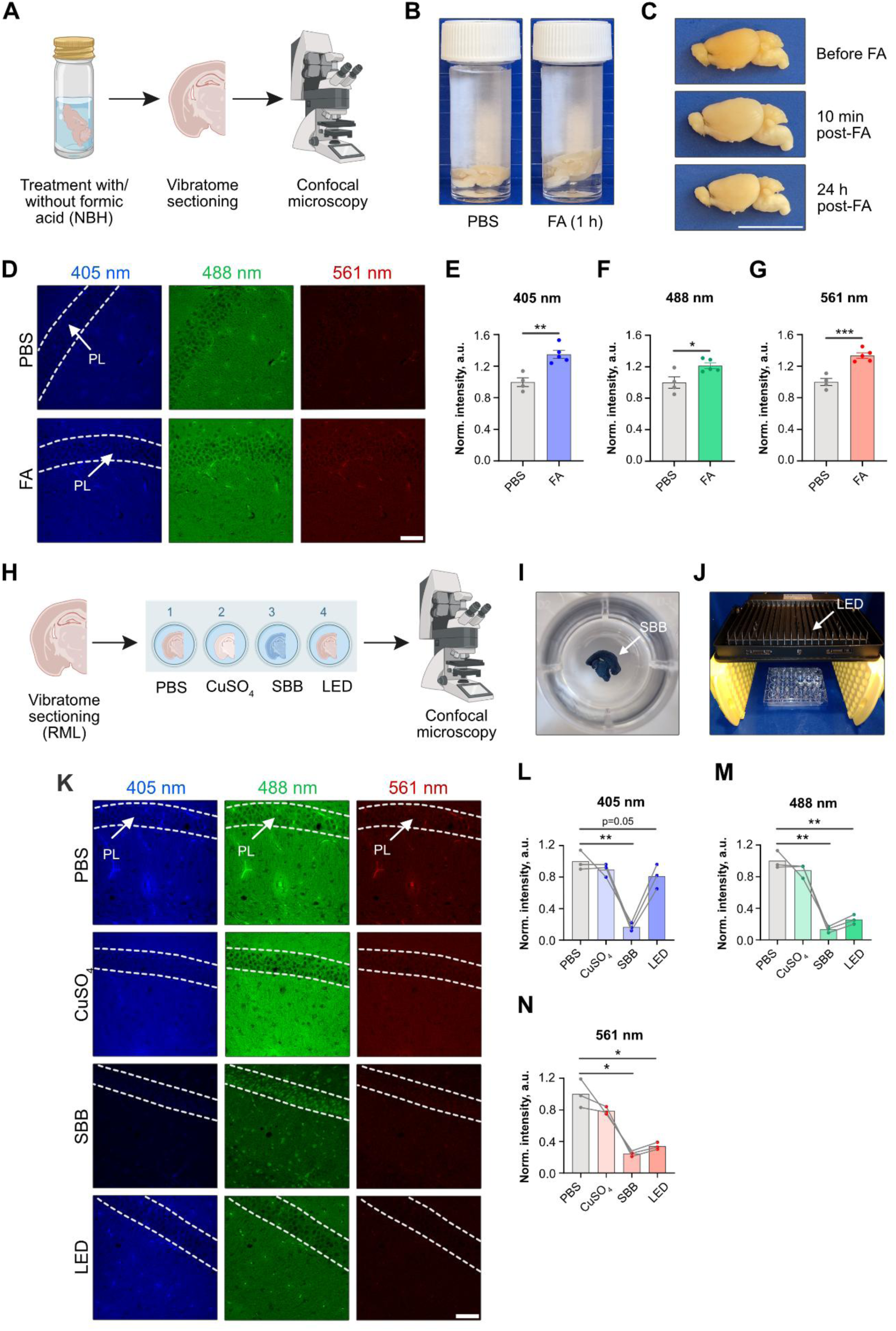
Application of autofluorescence quenching in prion-infected formic acid-treated mouse brains. **A)** The experimental design for assessing the effect of formic acid (FA) on tissue autofluorescence involved incubating hemispheres of NBH-injected mice with phosphate buffer saline (PBS, control) or FA, followed by vibratome sectioning and confocal microscopy of unstained samples. **B)** Photographs of mouse brains treated with PBS or FA for 1 h. **C)** Comparison of the same brain hemisphere before incubation with FA and at 10 min or 24 h post-treatment. **D)** Representative images of autofluorescence at 405, 488, and 561 nm in the hippocampal CA1 area. **E-G)** Plots of normalised tissue autofluorescence intensity at 405 nm (E), 488 nm (F), and 561 nm (G). **H)** The experimental scheme for evaluating the effectiveness of autofluorescence quenching approaches using free-floating vibratome-cut brain sections from RML scrapie prion-injected mice with PBS (control), copper sulfate (CuSO₄), Sudan Black B (SBB), or light-emitting diode (LED) illumination, followed by confocal microscopy of unstained samples. **I)** Photograph of a brain section after treatment with SBB, and **J)** setup for illuminating brain sections with an LED lamp. **K)** Representative images of autofluorescence at 405, 488, and 561 nm in the hippocampal CA1 area following different treatments. **L-N)** Plots of normalised tissue autofluorescence intensity at 405 nm (L), 488 nm (M), and 561 nm (N) under treatment conditions. Abbreviations: PL, pyramidal layer. Significance is indicated as p < 0.05 (*), p < 0.01 (**), p < 0.001 (***). Data is presented as mean ± SEM. The scale bars shown represent 10 mm (C) and 50 µm (D, K).

**Figure S2.**
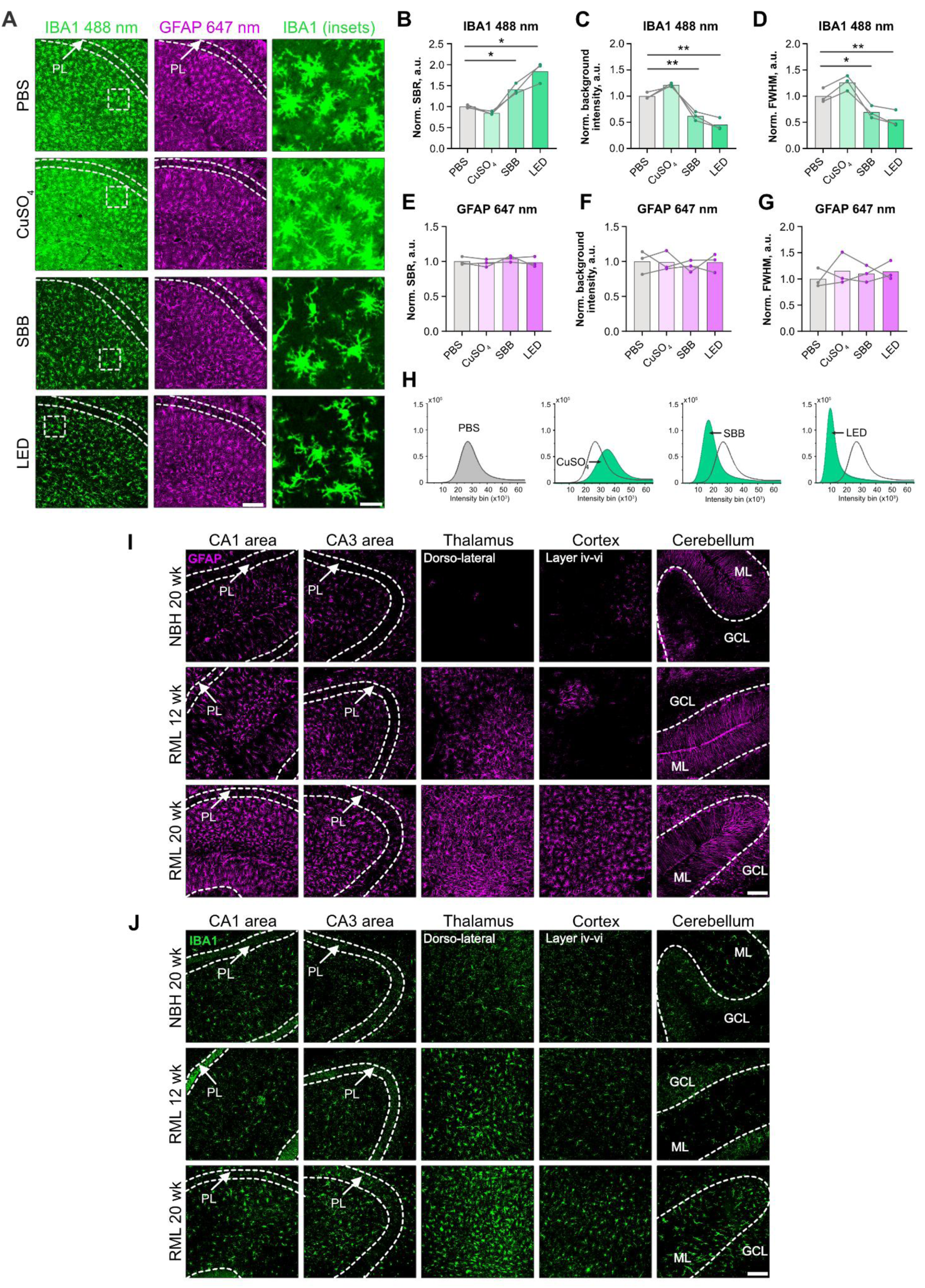
Immunofluorescent staining for glial markers in mouse brains following background quenching procedure. **A)** Representative images showing background quenching in the hippocampal CA1 area of RML scrapie-injected mice, labeled for ionised calcium-binding adaptor molecule 1 (IBA1; 488 nm, green) and glial fibrillary acidic protein (GFAP; 647 nm, magenta), after treatment with PBS (control), CuSO_4_, SBB, or LED illumination. The last column shows magnified views of IBA1⁺ microglia paired with dotted ROIs in the first column. **B-D)** Normalised signal-to-background ratio (B), background intensity (C), and full width at half maximum (FWHM) of intensity histograms for IBA1 at 488 nm (D). **E-G)** Normalised signal-to-background ratio (E), background intensity (F), and FWHM of intensity histograms for GFAP at 647 nm (G). **H)** Example intensity histograms of IBA1 staining following each treatment at 488 nm. **I-J)** Representative single-channel images of GFAP (I) and IBA1 (J) immunofluorescent staining in five brain ROIs of RML scrapie-infected mice after SBB treatment. Abbreviations: PL, pyramidal layer; ML, molecular layer; GCL, granule cell layer. Significance is indicated as p < 0.05 (*), p < 0.01 (**). Data is presented as mean ± SEM. The scale bars shown represent 100 µm (A, GFAP 647 nm; I and J) and 20 µm (A, IBA1 insets).

**Figure S3.**
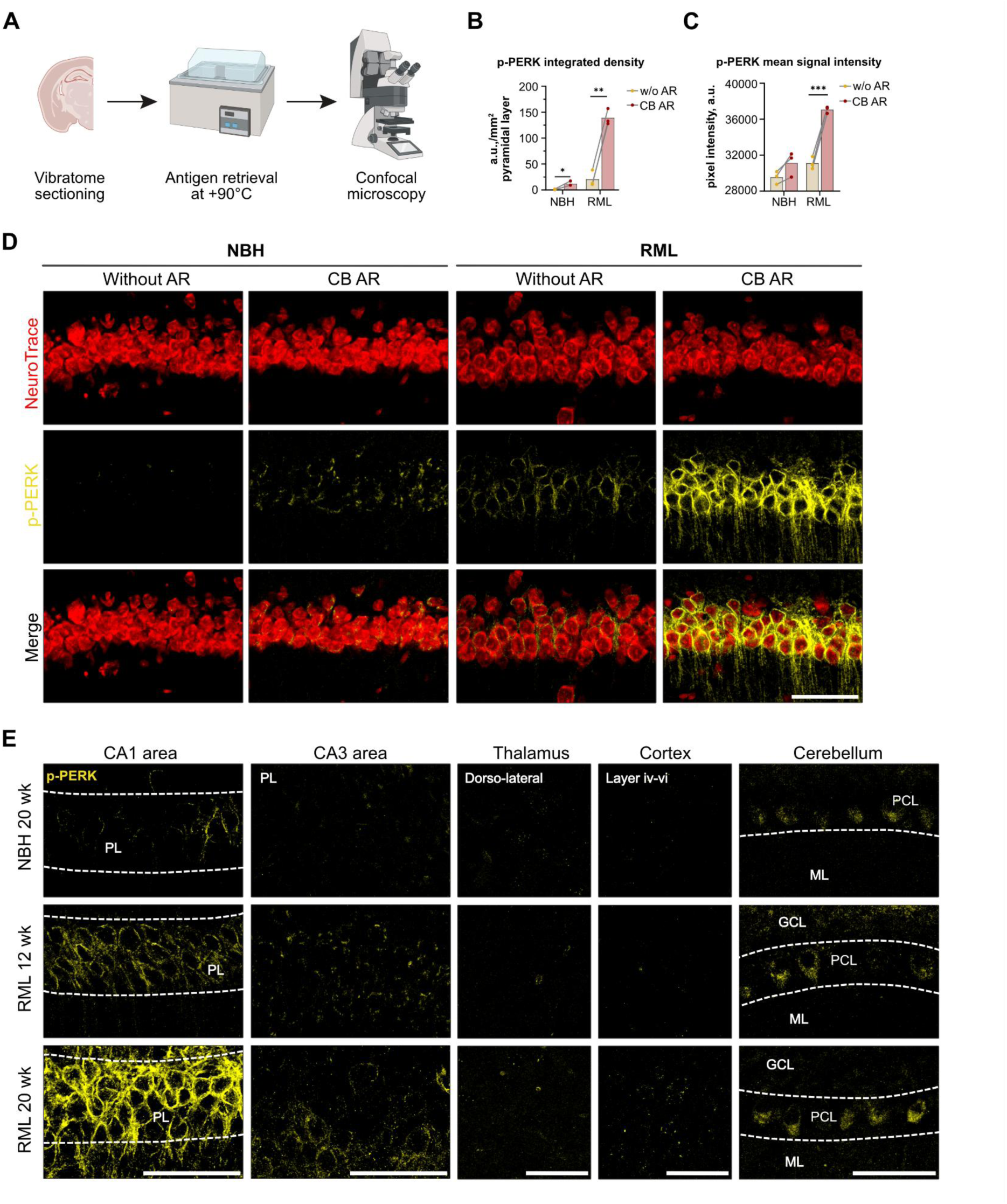
p-PERK immunofluorescence in the mouse brains with antigen retrieval. **A)** The experimental design for assessing the impact of heat-induced antigen retrieval on the fluorescent signal of p-PERK staining included incubating vibratome brain sections from NBH and RML mice in PBS at room temperature (control) or in citrate buffer on a water bath (CB AR), followed by immunohistochemistry and confocal microscopy. **B,C)** p-PERK⁺ integrated density per mm² in the pyramidal layer of the CA1 area (B) and p-PERK mean signal intensity (C). **D)** Representative images of p-PERK immunofluorescence (yellow) in the CA1 area with and without antigen retrieval, co-stained with NeuroTrace 660 (red). **E)** Representative images of p-PERK immunofluorescent staining in five brain ROIs of prion-infected mice without counterstaining. Abbreviations: PL, pyramidal layer; ML, molecular layer; PCL, Purkinje cell layer; GCL, granule cell layer. Significance is indicated as p < 0.05 (*), p < 0.01 (**), p < 0.001 (***). Data is presented as mean ± SEM. The scale bars shown represent 50 µm.

**Figure S4.**
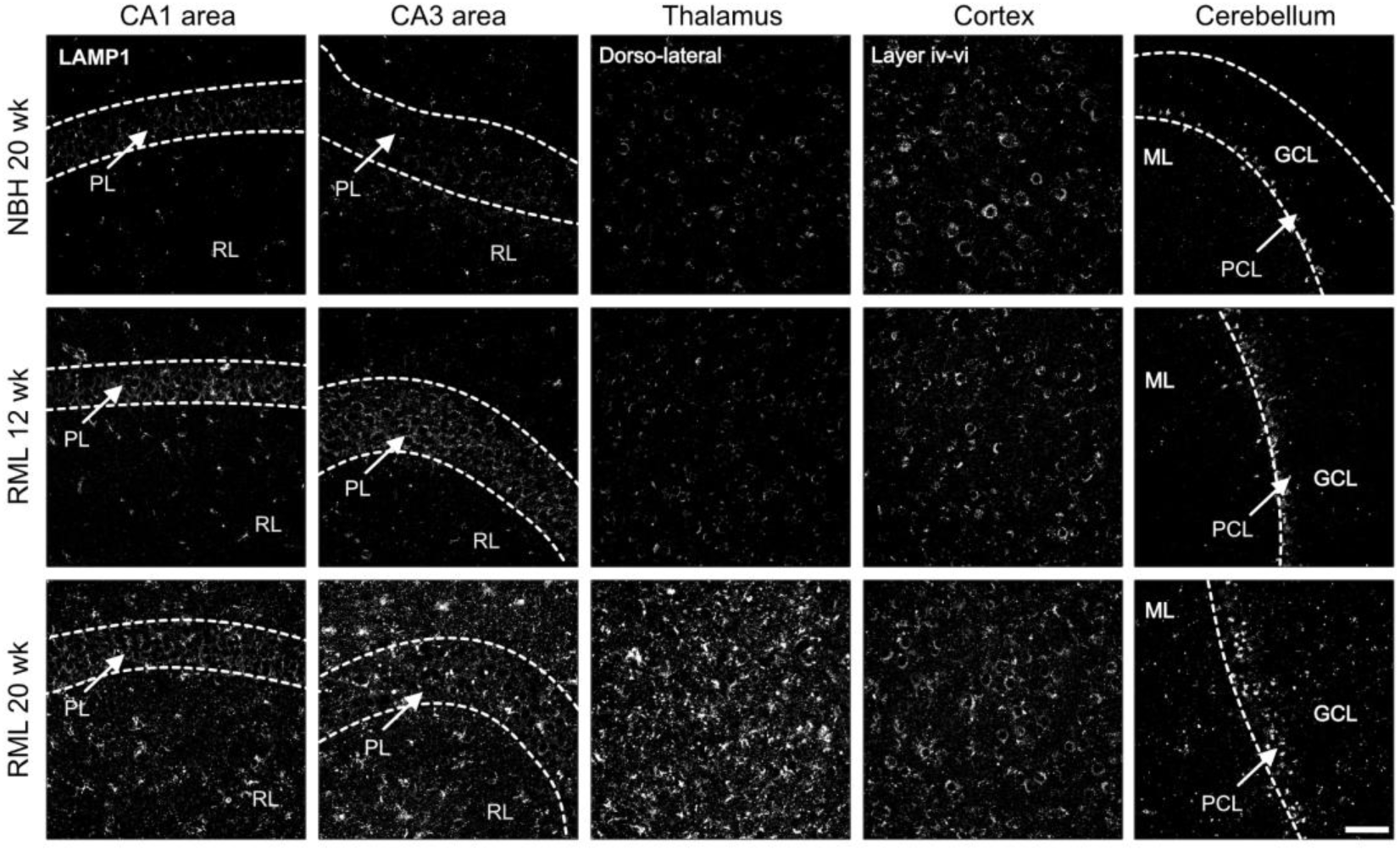
Representative images of LAMP1 immunofluorescent staining in five brain ROIs of RML scrapie-infected mice without counterstaining. Abbreviations: PL, pyramidal layer; RL, radial layer; ML, molecular layer; PCL, Purkinje cell layer; GCL, granule cell layer. The scale bar shown represents 50 µm.

